# The TREM2 targeting small molecule Sob-AM2 improves cognition independent of Aβ plaque alteration in 5xFAD mice

**DOI:** 10.64898/2026.07.31.741838

**Authors:** Lucas Kuhnau, Gisselle A. Jimenez, Wyatt Hack, Samantha Varada, Noah Gladen-Kolarsky, Sunwoo Kim, Finch Klein, Tapasree Banerji, Joseph F. Quinn, Thomas S. Scanlan, Nora E. Gray

## Abstract

**Background:** Inflammation is an early event that substantially influences Alzheimer’s disease (AD) pathogenesis, making it a compelling target for therapeutic intervention. Sob-AM2 is a brain-penetrating thyromimetic drug capable of inducing the expression of microglial cell surface receptor TREM2, which mediates the switch from pro-inflammatory to a more restorative microglial state.

**Objective:** Evaluate the effects of Sob-AM2 on cognition, AD pathology and microglial activity in the 5xFAD mouse model of amyloid-beta (Aβ) accumulation.

**Methods:** Seven-month-old 5xFAD mice and their wild-type littermates were administered Sob-AM2 subcutaneously three times per week for 12 weeks. In the last two weeks of treatment mice underwent behavioral tests to assess cognition and monitor for off-target mobility effects. At the end of treatment, brain tissue was harvested for gene and protein expression analyses.

**Results:** Sob-AM2 treatment increased TREM2 expression in the brains of 5xFAD mice. This was accompanied by an improvement in both spatial and associative memory as well as an increase in the expression of synaptic genes synaptophysin and PSD-95. No significant changes were detected in Aβ plaque burden or the expression of microglial activation marker Iba1 in Sob-AM2 treated animals, however, the expression of the phagocytic marker CD68 was significantly increased in the hippocampus, but not the cortex, in Sob-AM2 treated 5xFAD mice.

**Conclusion:** These results suggest that the cognitive-enhancing effects of Sob-AM2 are not the result of reduced overall plaque burden. Future work is needed to further investigate the neuroprotective mechanism of Sob-AM2 and how it may be affecting microglial phenotypes.

## 1. Background

Alzheimer’s disease (AD) currently affects over 7 million Americans aged 65 or older, and that number is expected to rise in parallel with the aging population [1]. Individuals living with AD experience profound impairments in cognition, deeply impacting their quality of life and necessitating extensive assistance from caregivers. The pathological hallmarks of AD include the accumulation of beta-amyloid (Aβ) plaques and neurofibrillary tau tangles, which are accompanied by progressive neuronal loss, inflammation, and eventual brain atrophy [2].

Although there is no cure for AD, efforts to intervene on disease progression have primarily focused on clearing the hallmark Aβ plaques. Despite the relative success of some of these amyloid-targeting therapies [3], their clinical benefit remains limited, as these treatments are effective only in select patient populations and when administered early in the disease course [4]. As a result, there has been growing interest in targeting other contributory cellular mechanisms that contribute to AD pathogenesis.

One such contributor is neuroinflammation, which has been shown to arise in the early stages of AD preceding clinical signs of cognitive impairment [5]. Neuroinflammation is mediated largely by microglia, the resident immune cells of the central nervous system (CNS). Microglia play essential roles under homeostatic conditions, including synaptic pruning, debris clearance, response to acute inflammatory signals, and removal of damaged or apoptotic neurons [6]. Even during the early stages of AD, microglia remain protective and work to clear Aβ plaques. However, the chronic activation associated with AD pathology leads microglia to adopt a disease-associated microglial (DAM) phenotype [7]. In this context, they can engulf healthy synapses, fail to clear Aβ plaques, and release pro-inflammatory cytokines that can damage healthy neurons [8]. Consistent with this, activated microglia are found around Aβ plaques and have been shown to release pro-inflammatory cytokines such as tumor necrosis factor alpha (TNF-α), interferon-gamma (IFN-y) or the interleukin (IL) like cytokines IL1-B and IL18 [9]. Together, these processes contribute to a sustained inflammatory environment in the brain that exacerbates neuronal dysfunction and degeneration.

Regulating microglial activity may offer a means of dampening AD-associated neuroinflammation. One promising genetic target for this approach to yield this regulation is triggering receptor expressed on myeloid cells 2 (TREM2), a transmembrane receptor expressed on both microglia and macrophages. TREM2, a member of the immunoglobulin superfamily, signals through the intracellular adaptor protein DAP12 to regulate microglial functions like inflammatory cytokine release and phagocytosis [10]. In fact, phagocytic activity in Aβ clearance is thought to be dependent on TREM2 activation [11].

TREM2 has been identified as a significant genetic risk factor of AD [12]. In particular, the missense mutation R47H has been associated with a nearly three-fold increase in AD risk [13]. A recent study using human microglia cells showed that TREM2 loss and haploinsufficiency impair microglial responses to Aβ deposition and enhance Aβ-induced local neurotoxicity [14]. Conversely, TREM2 overexpression in AD mouse models has been shown to reduce levels of proinflammatory transcripts and ameliorate AD-related neuropathology [15]. Together, these findings position TREM2 as a key regulator of microglial function and disease progression in AD and have motivated efforts to therapeutically target this pathway.

One strategy to increase TREM2 expression is through transcriptional induction. Our group recently reported a thyroid hormone receptor (THR) binding site in the promoter region of TREM2 that can be induced using compounds that bind and activate THRs [16]. One such compound is sobetirome, a small molecule mimetic of the biologically active thyroid hormone triiodothyronine (T3). Sobetirome acts as a selective agonist for THR-β, a receptor commonly targeted for metabolic therapies due to its predominant expression in the liver and pituitary gland [17]. The amide prodrug of sobetirome, Sob-AM2, was developed to enhance CNS delivery, and was found to increase sobetirome brain exposure while minimizing peripheral tissue exposure, thereby reducing potential systemic side effects [18]. Sob-AM2 has been shown to increase TREM2 expression and reduce neuroinflammation and disease severity in an experimental autoimmune encephalomyelitis (EAE) model of multiple sclerosis [19, 20].

Given the strong link between neuroinflammation, TREM2 and AD disease progression, these findings suggest that Sob-AM2 could be therapeutically beneficial in AD models as well. This study aims to evaluate Sob-AM2 in the context of AD using the 5xFAD mouse model of Aβ accumulation to assess its effects on cognition, Aβ accumulation, microglia activity, and synaptic integrity.

## 2. Methods

### 2.1. Animals

Experiments were conducted in line with the NIH Guidelines for the care and use of laboratory animals and approved by the Institutional Animal Care and Use Committee (IACUC) of the Veteran’s Administration Portland Health Care System (VAPORHCS; ACORP #6155-23). 5xFAD transgenic mouse colonies were generated by breeding hemizygous males to non-carrier females obtained from The Jackson Laboratory (Bar Harbor, ME, USA). These mice overexpress human amyloid precursor protein (APP) and human presenilin 1 (PS1) due to five mutations associated with Familial Alzheimer’s Disease (FAD): the Swedish (K670N, M671L), Florida (I716V), and London (V717I) mutations in APP, and the M146L and L286V mutations in PS1 [21].

These transgenic mice develop beta-amyloid (Aβ) plaques around 2 months of age, with synaptic loss, elevated neuroinflammation, and spatial memory deficits evident by 4-5 months of age [21]. Animals were kept in a climate-controlled environment with a 12-hour light/dark cycle and provided with water and diet *ad libitum*.

### 2.2. Dosing regimen and behavioral schedule

WT and 5xFAD mice were randomly assigned to Sob-AM2 or control groups with 11-12 animals of each sex per group (Figure 1A and W1B). The sample size for the study was determined from the expected number of mice per group necessary to observe changes in Aβ pathology as well as CFR and OLM performance with 80% power assuming two-sided significance of 0.05, based on existing data from our group and others using 5xFAD mice [22–25]. We calculated that 10 mice per group was necessary to detect changes for those endpoints. In the study, we used groups of 12 to account for potential attrition.

**Figure 1:**
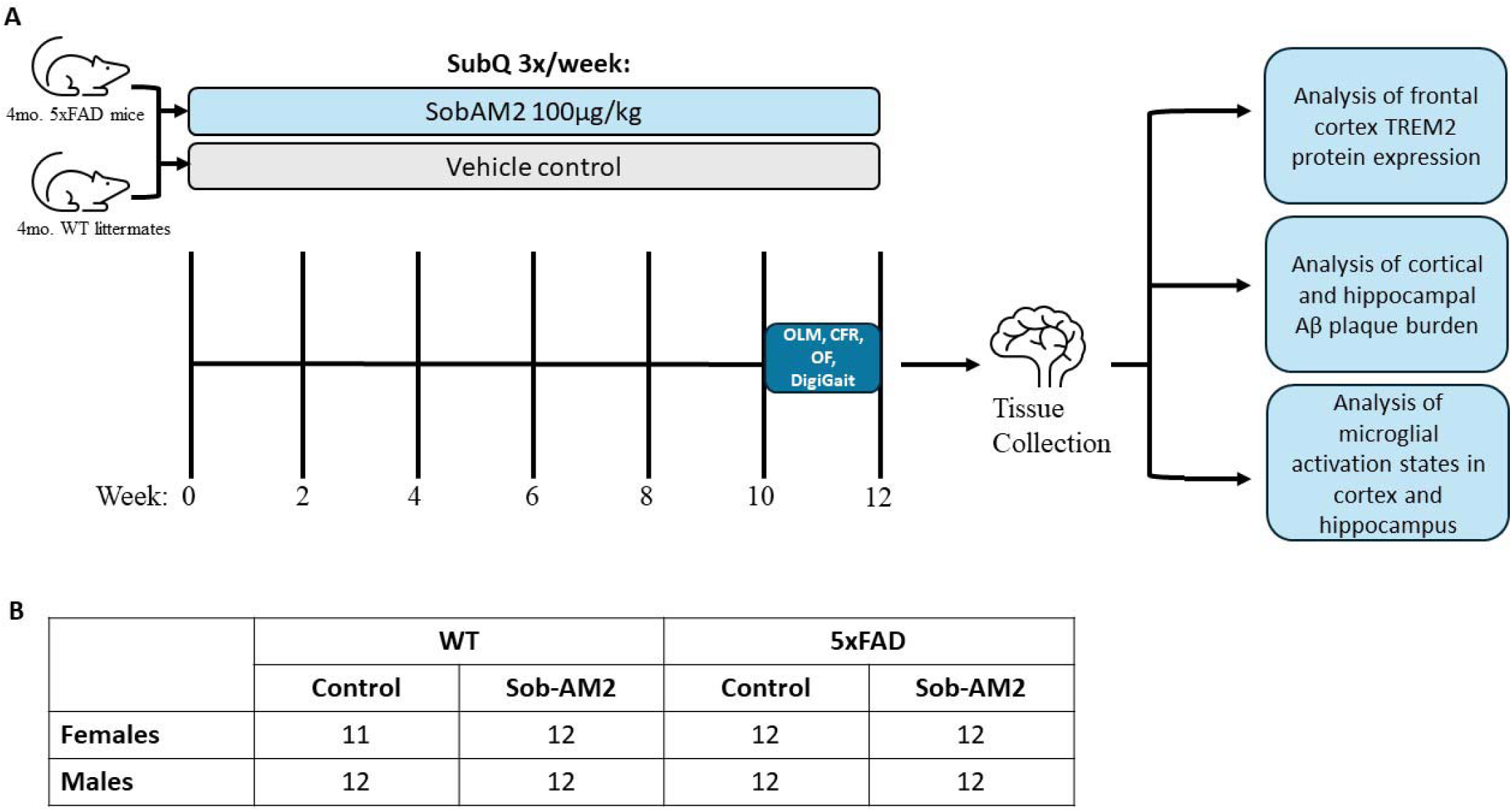
Experimental design. (A) Treatment and behavioral testing timeline. (B) Number of animals in each treatment group.

At 4 months of age, male and female 5xFAD mice and WT littermates began treatment with either 100 µg/kg of Sob-AM2 in 0.9% physiological saline or just 0.9% physiological saline as a vehicle control, administered subcutaneously 3 times a week for a total of 12 weeks. The chemical synthesis of Sob-AM2 was reported previously [26]. In the final 2 weeks of treatment, mice underwent cognitive testing. Female mice were housed 3-4 per cage, while male mice were either housed 3-4 to a cage or sometimes 1-2 per cage due to aggression. Body weight was monitored throughout the 12-week treatment period, and no significant differences were observed across treatment groups (Supplementary Figure 1).

In the final two weeks of treatment, animals underwent behavioral testing (described below). Tests were always conducted in the same order with Object Location Memory (OLM) test first, followed by the Open Field (OF) test, then Conditioned Fear Response (CFR) and finally DigiGait. Behavioral testing occurred only at the end of the treatment paradigm to avoid the confound of learning on test performance.

### 2.3. Open Field (OF)

The OF test is an assessment of overall mobility. Each mouse was placed into a square arena (38 cm × 38 cm × 64 cm high, constructed of white acrylonitrile butadiene styrene) for a 5-minute period. A camera mounted above the arena, interfaced with the AnyMaze video tracking system (Stoelting Co, Wood Dale, IL, USA) captured distance traveled (mm), which is a metric of overall mobility.

### 2.4. Object Location Memory (OLM)

The OLM is a hippocampal-dependent test that assesses spatial memory [27]. It is a four-day procedure (Figure 2). Days 1 and 2 each consist of a ten-minute session where mice are habituated to the testing chamber (39 cm x 39 cm x 39 cm). Training begins on the third day and involves three 10-minute sessions where mice interact with a pair of identical objects that are placed in a fixed location in the upper corners of the testing chamber. Each training session occurs approximately an hour after the previous training session. The training sessions are followed by two retention tests, which occur 2 and 24 hours after the final training session. In each 5-minute retention test, one of the objects remains in the same location as it was for training while the other object is moved to a novel location in the testing chamber for each test session. An overhead camera captures video of the animals throughout the test sessions and the time spent exploring the object in the novel location as well as the object in the familiar location were scored manually by an investigator blinded to treatment conditions using AnyMaze software version 7.48. To prevent the influence of inter-rater variability, all tests were scored by the same investigator. The time spent exploring the object in the novel location relative to the total time exploring both objects was expressed as a percentage. Increased time spent exploring the object in the novel location reflects improved spatial memory, based on the exploratory nature of mice [27]. The objects used were of similar height and width. Mice whose combined total exploration time for both objects was fewer than 5 seconds during a given retention trial were considered non-participating, and their data was excluded for that trial.

**Figure 2:**
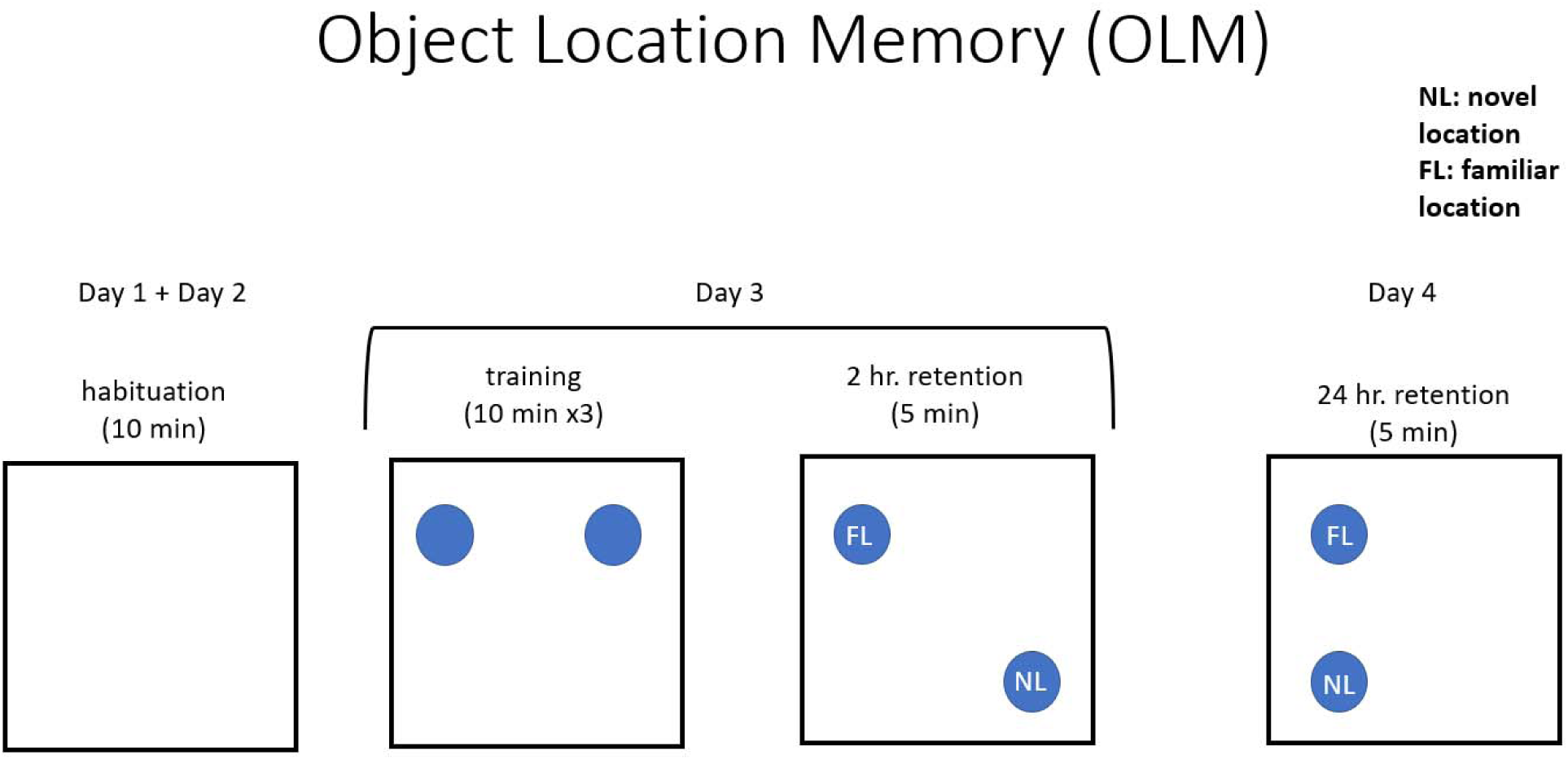
Diagram of OLM experimental set-up.

### 2.5. Conditioned Fear Response (CFR)

The CFR test assesses associative memory, and is mediated by inputs from the amygdala, hippocampus, and cortex [28]. It is a two-day procedure involving three phases: habituation, conditioning, and testing (Figure 3). The first day begins with habituation, where animals freely explore a chamber (16 x 16 x 12”) with a wire floor for five minutes. The conditioning phase is initiated immediately after habituation wherein the animal receives three 1-second shocks (0.5 mA), randomly distributed over a three-minute period. The testing phase occurs 24 hours after the conditioning phase, and involves the animal being re-introduced to the same chamber for two consecutive five-minute sessions without receiving any shocks. The amount of time spent freezing during each testing phase is automatically scored by AnyMaze software using the video feed from an overhead camera.

**Figure 3:**
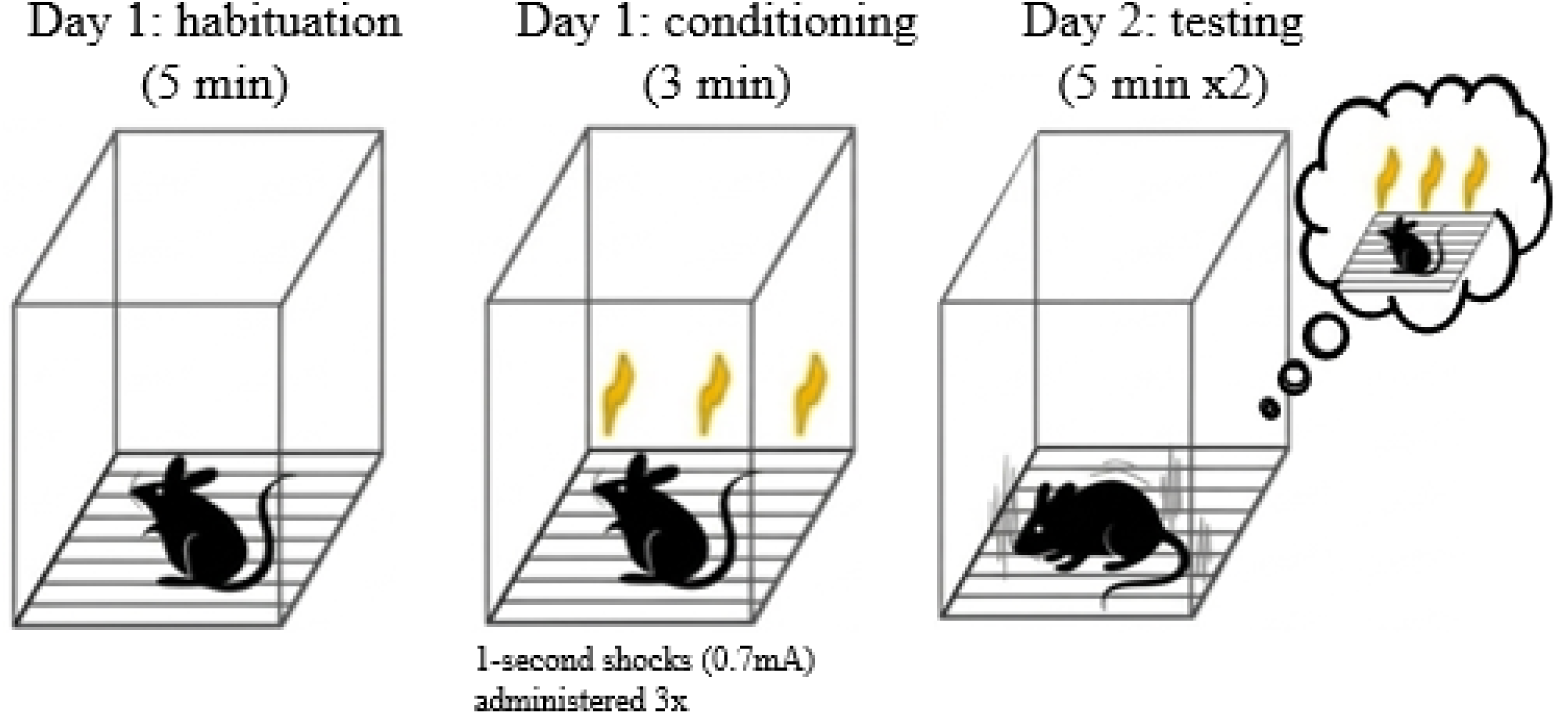
Diagram of CFR set-up.

The amount of time spent freezing during the habituation phase is then subtracted from each value to yield the change in freezing time from baseline and account for differences in overall activity between different strains of mice that could confound data interpretation. A greater increase in freezing compared to baseline reflects better contextual associative memory, based on the animal associating the context of the testing chamber with the shocks received in it [28].

### 2.6. DigiGait

The DigiGait apparatus (Mouse Specifics, Quincy, MA) allows users to analyze gait differences through recorded videos of mouse movement along a transparent treadmill belt taken from a ventral angle. Mice were first introduced to the treadmill area, which was bound by an acrylic compartment (5 cm in width, 25 cm in length). Mice were given one minute to habituate to the treadmill compartment, after which the treadmill motor was activated at a speed of 20 cm/second with a 15-degree incline. To acquire usable data, mice needed to move for 5-6 consecutive seconds without rearing, stopping, or jumping to provide an anticipated 16+ consecutive strides for data analysis. Videos were then analyzed by an individual, blinded to experimental conditions, using the DigiGait software (version 16) to validate automated identification of strides taken by each paw and extract data on gait metrics.

### 2.7. Euthanization and tissue collection

Following behavioral tests, mice were euthanized using inhalable isoflurane, cardiac puncture and cerebral dislocation. Brain samples were collected. The anterior 3 mm of bilateral prefrontal cortex was dissected along with the cerebellum and brainstem. The lateral hemispheres were then separated, and the remaining left hemisphere was subdissected and frozen. The right hemisphere of each sample was immersed in 4% paraformaldehyde (TCI Chemicals) diluted in phosphate-buffered saline (PBS) for fixation for 24 hours. Following fixation, tissues were incubated in PBS for 24h and then cryoprotected in a graded sucrose series (15% followed by 30% sucrose in PBS) for 24h each and then frozen at -80°C until sectioning. Samples were sectioned at 40 µm thickness using a freezing microtome. The contralateral cortex was subdissected and frozen.

### 2.8. Immunohistochemistry (IHC) analysis

Sections of similar anatomical depth were placed in quenching solution (30% methanol, 10% hydrogen peroxide, and 10% Tris-HCl buffered saline) for 2 minutes to inactivate endogenous enzymes and then incubated in blocking buffer (2% bovine serum albumin, 10% donor horse serum, 0.5% Triton X-100, 10% Tris-HCl buffered saline) for 3 hours to prevent non-specific antibody binding. Sections were then incubated overnight with primary antibodies diluted in 1x phosphate-buffered saline (PBS): anti-Aβ polyclonal antibody (Kerafast, EB2001), anti-Iba1 (FujiFilm,019-19741), and anti-CD68 (Invitrogen,14-0681-82) at concentrations of 1:1000, 1:2500, and 1:300, respectively. The next day, sections were incubated in biotinylated IgG secondary antibodies (Aβ: anti-mouse at 1:1000; Iba1: anti-rabbit at 1:1000; CD68: anti-rat at 1:300) for 2 hours. Signal detection was performed using a diaminobenzidine (DAB) tablet set as previously described [29]. Sections were mounted on UltraSlips cover glass slides (GC2450-ACS) and scanned using PrimeHisto XE (Pacific Image Electronics, Torrance CA, USA). ImageJ software (Rasband, W.S., ImageJ, U.S. National Institutes of Health, Bethesda, Maryland, USA, https://imagej.nih.gov/ij/, 1997–2018) was used to quantify images. Images were converted to 8-bit grayscale, and a human user, blinded to the treatment conditions, employed the polygon tool to encompass the hippocampus or cortex and the total area of each brain was recorded. The threshold was adjusted to remove ubiquitous background staining, highlighting only areas of intense staining. Staining was quantified in three coronal sections representing regions of anterior, middle and posterior hippocampus and cortex from each right hemisphere sample. Hippocampal and cortical areas were traced using a computerized stage and stereo investigator software. Quantification of Aβ plaques as well as Iba1 and CD68 staining was expressed as a percentage of the cortical or hippocampal area (cm^2^) that was occupied by detectable immunoreactive staining. Mean values for each parameter were calculated using at least three sections per animal.

### 2.9. TREM2 quantification

Protein was extracted from each sample using the frontal cortex. Briefly, half of the frontal cortex was homogenized in TRI (Thermo Fisher Scientific, 15596026) and BAN reagents (Molecular research Center, BN191). After centrifugation at 12,000 RPM at 4°C, the aqueous phase was carefully separated from the protein phase. The protein was subsequently washed five times with centrifugation between washes to ensure purity. The resulting protein pellet was resuspended in 250 ul SDS containing 1% protease and phosphatase inhibitor (Thermo Fisher Scientific, 1861282). Protein concentrations of TREM2 were determined by ELISA (Reddot Biotech, RD-TREM2-Mu). A bicinchoninic acid (BCA) assay was used to determine total protein concentration of our samples and 80ug/ul of protein per well was used for the ELISA assay which was performed according to the manufacturer’s instructions.

### 2.10. Gene expression analysis

RNA was extracted from the aqueous phase of the frontal cortex Tri Reagent extraction described above per the manufacturer’s instructions. RNA was then reversed transcribed to cDNA using the SuperScript III First-Strand Synthesis kit (Thermo Fisher Scientific, 18080051) per the manufacturer’s instructions. Relative gene expression was assessed using TaqMan Fast Advanced Master Mix (Applied Biosystems by Thermo Fisher Scientific, 4444557) and commercially available TaqMan primers (Thermo Fisher Scientific) for synaptophysin (Mm00436850_m1), post-synaptic density protein 95 (PSD-95; Mm00492193_m1), and glyceraldehyde- 3phosphate dehydrogenase (GAPDH; Hs02758991_g1). Quantitative PCR (qPCR) was performed on the QuantStudio 3 Real-Time PCR System (Applied Biosystems by Thermo Fisher Scientific, A28137) and analyzed using the delta-delta Ct method.

### 2.11. Statistics

Graphs reflect groups averages with error bars indicating the standard error of the mean. Statistical differences were determined by ANOVA followed by Tukey pairwise comparisons or by T-test when only two groups were compared. All analyses were performed using GraphPad Prism (Version 11.0.0). Data is presented with males and females together as there was no significant interaction between treatment and sex.

## 3. Results

### 3.1. Sob-AM2 significantly increases TREM2 expression in the frontal cortex of 5xFAD mice

Sob-AM2 treatment significantly increased cortical TREM2 expression in 5xFAD animals compared to vehicle-treated 5xFAD mice (Figure 4). Vehicle-treated 5xFAD mice exhibited a trend towards lower expression of TREM2 in the frontal cortex compared to WT animals.

**Figure 4:**
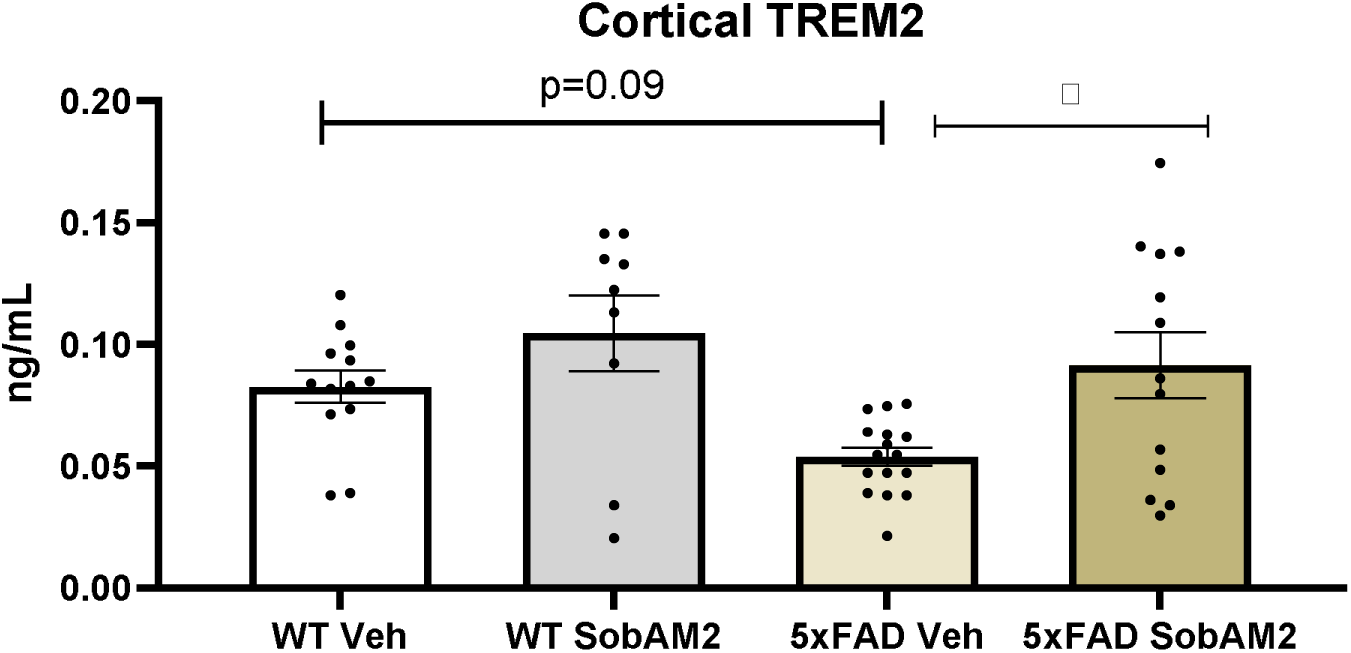
Sob-AM2 increases TREM2 expression in the frontal cortex of 5xFAD mice. Sob-AM2 treatment increased TREM2 expression in both WT and 5xFAD mice. A significant increase was observed in Sob-AM2 treated 5xFAD mice compared to their vehicle-treated controls. *p<0.05, WT Veh n= 22, WT SobAM2 n= 20, 5xFAD Veh n= 18, 5xFAD SobAM2 n= 19, F (3,47) =5.1.

### 3.2. Sob-AM2 improves spatial memory in 5xFAD mice

Spatial memory was assessed using the OLM test at 2 and 24 hours after the conclusion of task training. At both points, vehicle-treated 5xFAD mice spent a significantly lower percentage of their exploration time investigating the object in the novel location relative to their WT littermates (Figure 5A and B), indicating impaired spatial memory. However, Sob-AM2 treatment significantly improved OLM performance in 5xFAD mice at both 2h (Figure 5A) and 24h (Figure 5B) to levels similar to WT animals. Sob-AM2 treatment did not affect OLM performance for WT animals.

**Figure 5:**
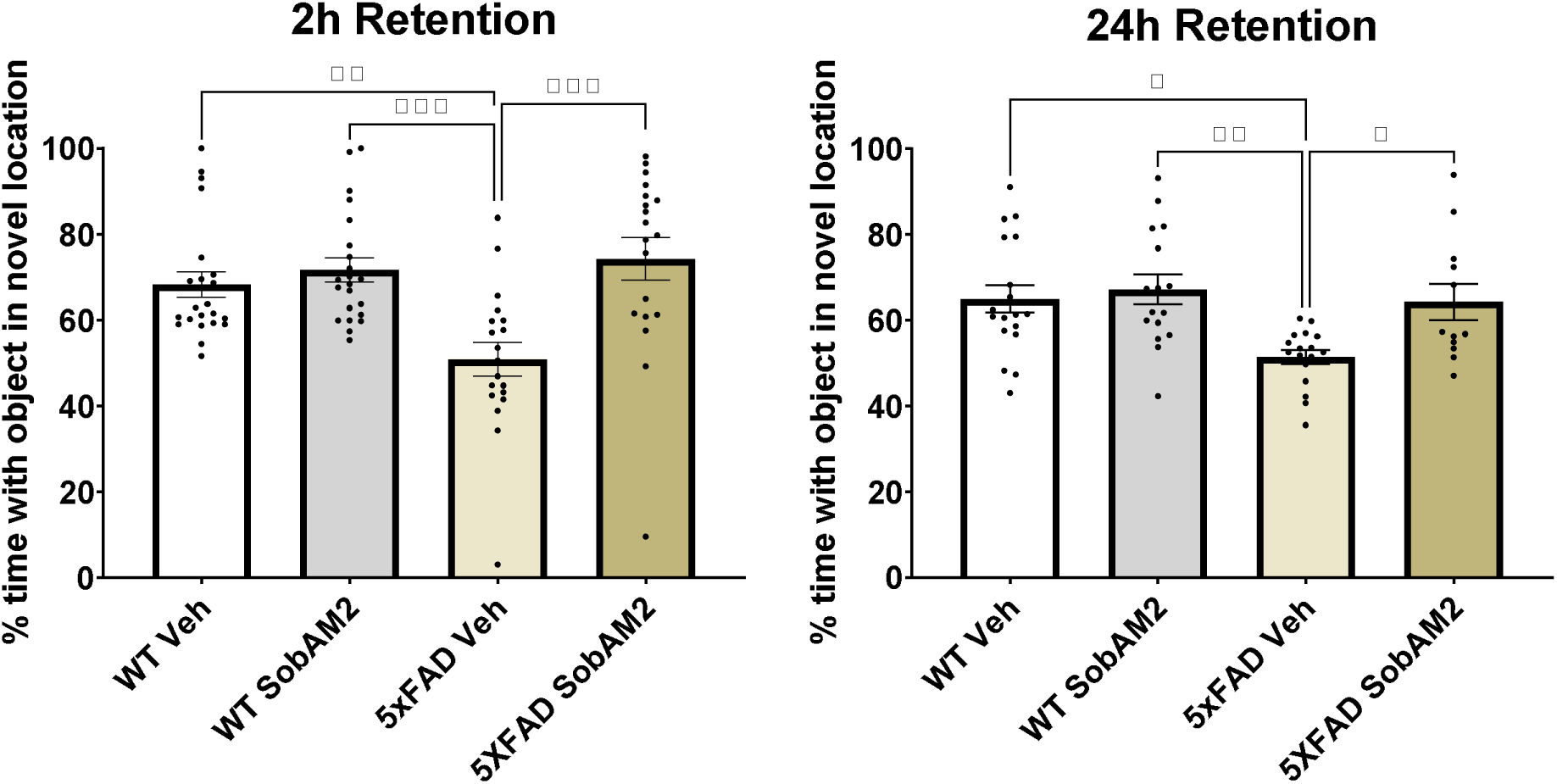
Sob-AM2 improves spatial memory in 5xFAD mice. Sob-AM2 treatment attenuated deficits in OLM performance in 5xFAD mice at both the 2h (A) and 24h (B) tests. *p<0.05, **p<0.01, ***p<0.001, 2H: WT Veh n= 23, WT SobAM2 n= 24, 5xFAD Veh n= 16, 5xFAD SobAM2 n= 21, F (3,78) =7.901. 24HR: WT Veh n= 19, WT SobAM2 n= 17, 5xFAD Veh n= 15, 5xFAD SobAM2 n= 17, F (3,59) =5.549.

### 3.3. Sob-AM2 treatment improves associative memory in 5xFAD mice

The CFR test was used to assess associative memory, In the vehicle treated group, 5xFAD mice displayed a significant reduction in freezing compared to vehicle-treated WT mice, indicating impaired memory. Sob-AM2 treatment significantly improved CFR performance in 5xFAD mice but had no effect on performance in WT animals (Figure 6).

**Figure 6:**
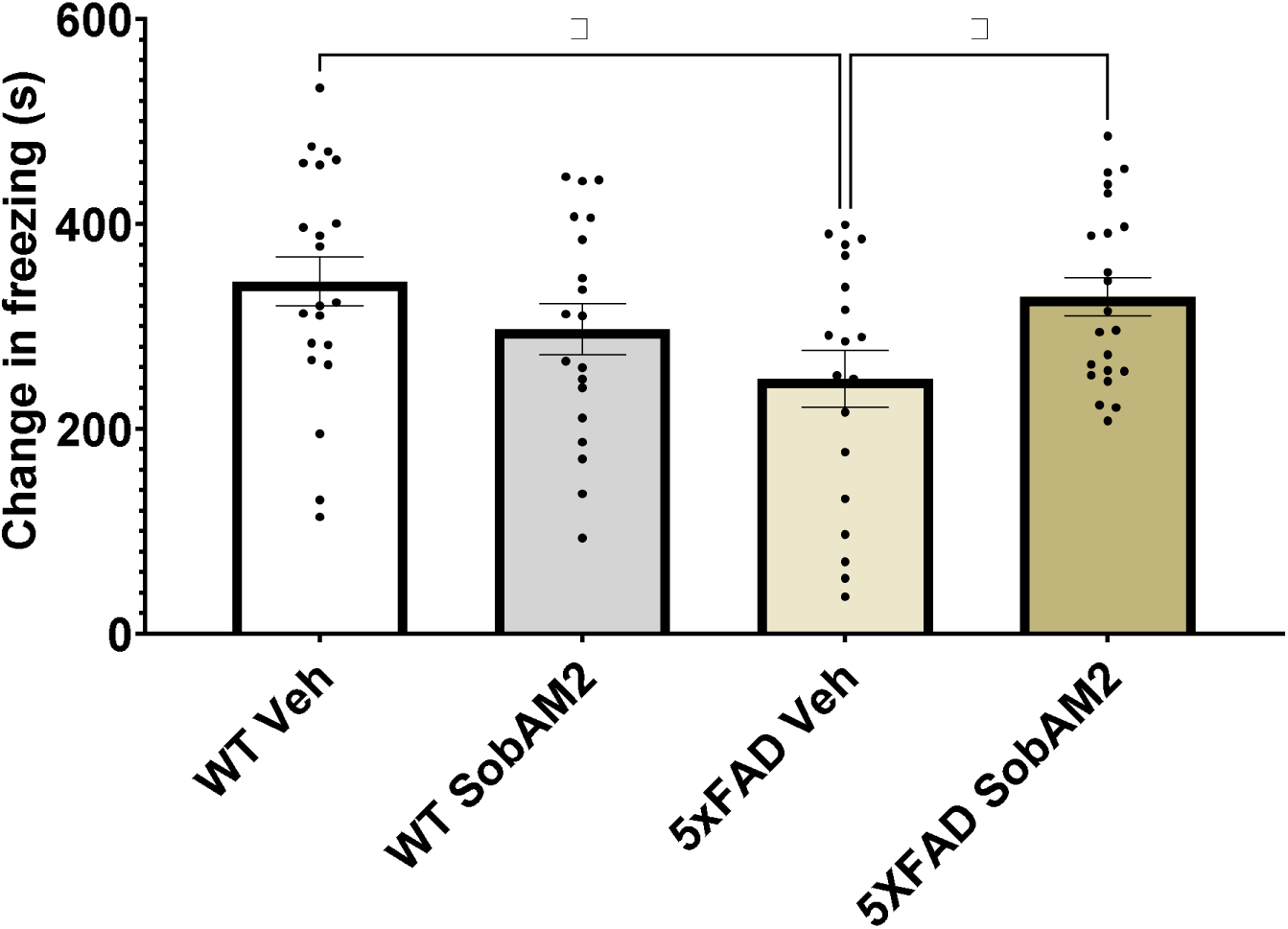
Sob-AM2 treatment improves associative memory in 5xFAD mice. A significant improvement in CFR performance is seen in Sob-AM2-treated 5xFAD mice over their vehicle-treated 5xFAD counterparts. *p<0.05, WT Veh n= 24, WT SobAM2 n= 24, 5xFAD Veh n= 22, 5xFAD SobAM2 n= 24, F (3,78) =3.07.

### 3.4. Sob-AM2 does not alter mobility in 5xFAD mice

Because Sob-AM2 is a thyromimetic compound, and thyroid hormones regulate metabolism and muscle function, which are critical for mobility [1], overall motor function was evaluated using the Open Field test. Sob- AM2 administration did not significantly alter the distance traveled in the arena compared to vehicle-treated animals in either WT or 5xFAD mice. Although 5xFAD mice exhibited reduced locomotor activity relative to their WT littermates, this difference was not statistically significant (Figure 7).

**Figure 7:**
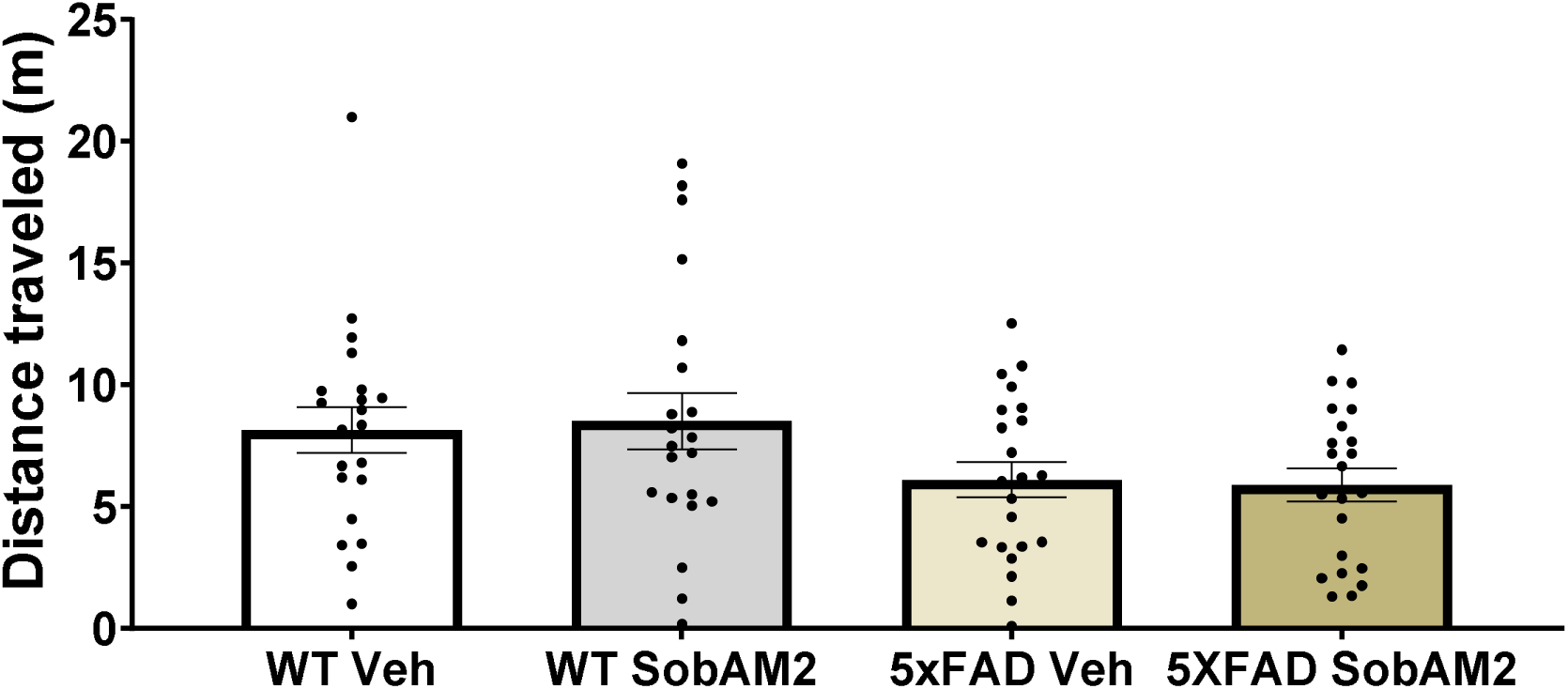
Sob-AM2 does not alter mobility in 5xFAD mice. 5xFAD mice exhibited a non-significant trend towards a lesser distance traveled but this was unaffected by Sob-AM2 treatment. Sob-AM2 likewise did not significantly alter distance traveled in WT mice. WT Veh n= 21, WT SobAM2 n= 23, 5xFAD Veh n= 23, 5xFAD SobAM2 n= 24, F (3,82) =2.334.

Additional metrics of gait were quantified using the DigiGait apparatus. Of the 32 metrics of gait quantified there were no significant differences found to be altered by Sob-AM2 in the 5xFAD mice with the exception of slight change in absolute paw angle for the left hind paw (Supplementary Table 1) which actually brought the value closer to what was observed in the WT vehicle treated animals.

### 3.5. Sob-AM2 increases synaptic gene expression in both 5xFAD and WT mice

Sob-AM2 treatment resulted in a significant increase in the expression of synaptophysin (Figure 8A) and PSD- 95 (Figure 8B) in the frontal cortex of 5xFAD mice. A non-significant decrease in the expression of synaptophysin (WT Veh=1.0 +/- 0.27, 5xFAD Veh=0.74 +/- 0.13) and PSD-95: (WT Veh=1.0 +/- 0.26, 5xFAD Veh=0.82 +/- 0.19). was observed in vehicle-treated 5xFAD mice relative to vehicle treated WT mice

**Figure 8:**
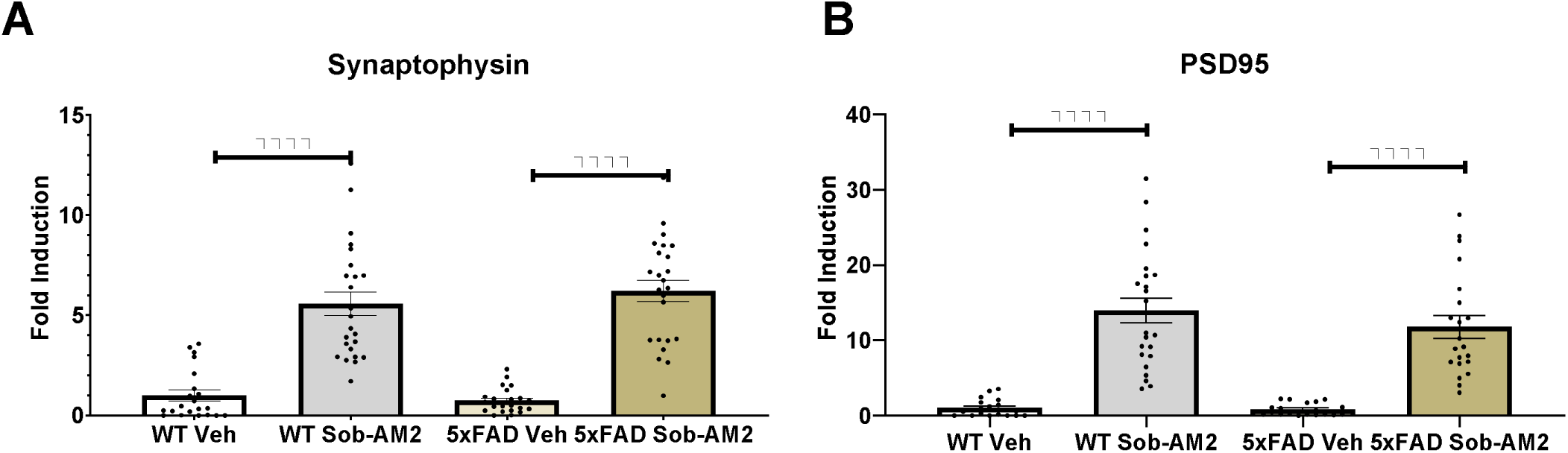
Sob-AM2 treatment significantly increases synaptic integrity across genotypes. Sob-AM2 treatment significantly increased the expression of synaptophysin (A) and PSD-95 (B) in both WT and 5xFAD mice compared to their vehicle-treated counterparts. ***p<0.0001. Syn: WT Veh n=21, WT Sob-AM2 n=24, 5xFAD Veh n=22, 5xFAD Sob-AM2 n=24, F (3,87) =42.76. PSD-95: WT Veh n=19, WT Sob-AM2 n=23, 5xFAD Veh n=19, 5xFAD Sob-AM2 n=21, F (3,78) =32.64.

### 3.6. Sob-AM2 treatment does not alter A**β** plaque burden in 5xFAD mice

There was no change in Aβ plaque abundance in 5xFAD mice following Sob-AM2 treatment (Figure 9A . Plaque burden was similar in both the hippocampus (Figure 9B) and the cortex (Figure 9C) in Sob-AM2 treated 5xFAD mice compared to vehicle-treated 5xFAD mice.

**Figure 9:**
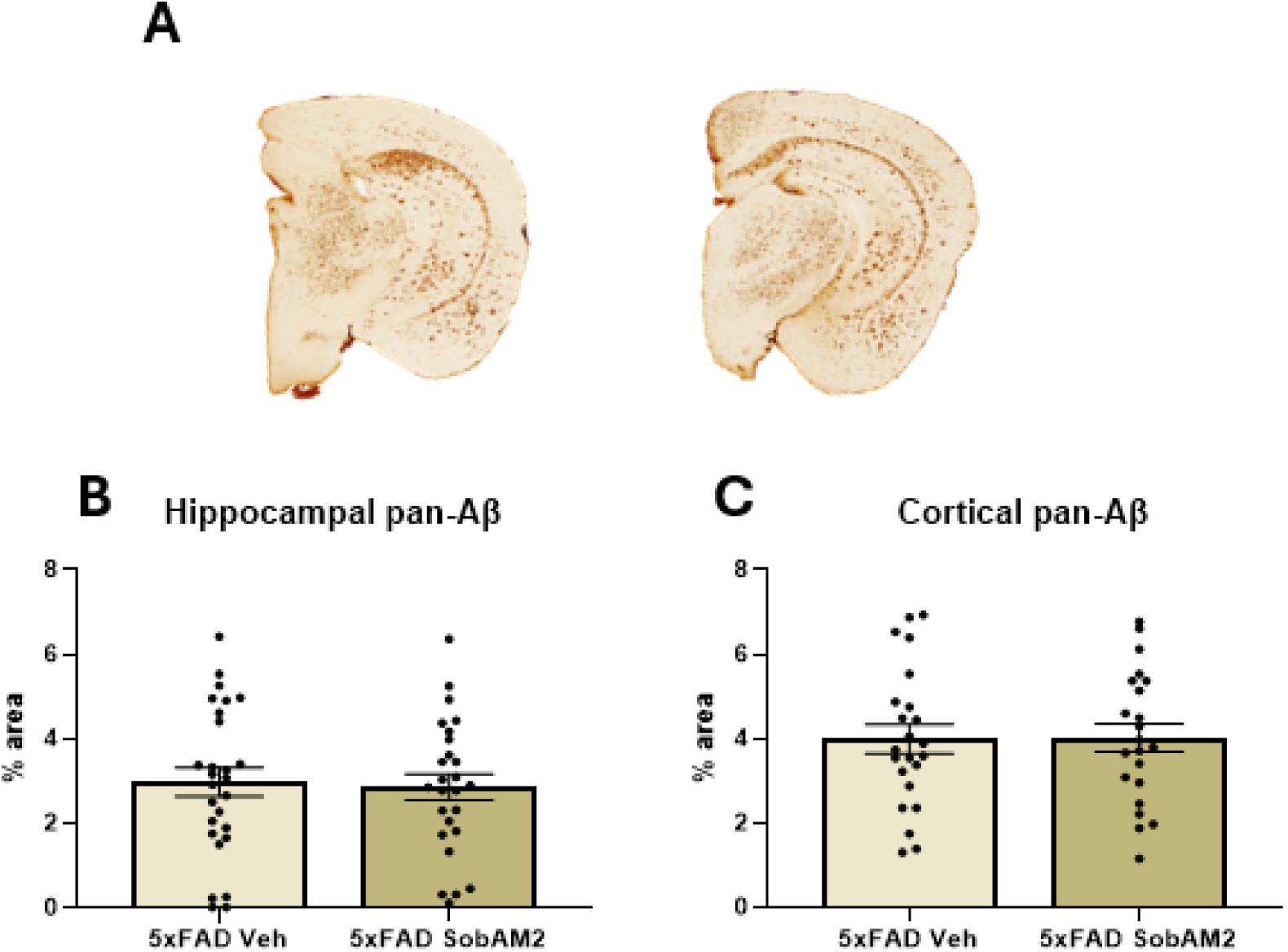
Sob-AM2 treatment does not alter. **A**β plaque burden in 5xFAD mice. (A) Representative images of AB staining in 5xFAD mice. Sob-AM2 treatment did not alter the overall AB plaque burden in either the (B) hippocampus, 5xFAD Veh n= 27, 5xFAD Sob-AM2 n= 26 or (C) cortex, 5xFAD Veh n= 23, 5xFAD Sob-AM2 n= 22.

### 3.7. Sob-AM2 does not alter microglial activation in 5xFAD mice

Iba1 protein expression was quantified as a marker of microglial activation. WT mice expressed little to no Iba1, while 5xFAD showed significantly elevated levels, regardless of treatment (Figure 10A). There was no effect of Sob-AM2 treatment on Iba1 expression in either the hippocampus (Figure 10B) or cortex (Figure 10C).

**Figure 10:**
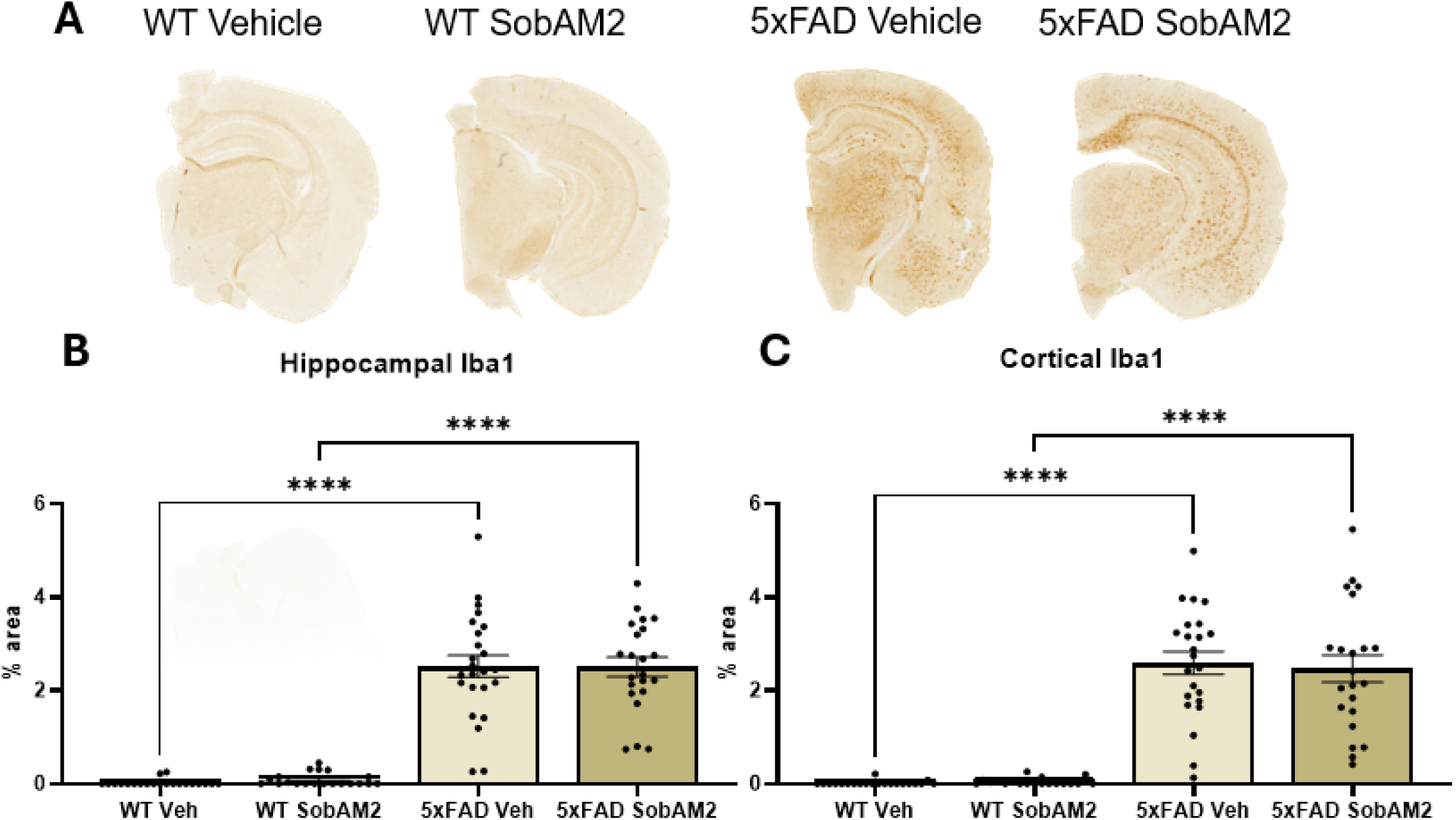
Sob-AM2 does not alter microglial activation in 5xFAD mice. (A) Representative images of Iba1 staining in WT and 5xFAD mice. 5xFAD mice displayed significantly higher Iba1 expression compared to their littermate controls. Sob-AM2 treatment showed a trend toward reduced Iba1 expression; however, this reduction did not reach statistical significance in either the (B) hippocampus or (C) cortex. Hippocampus: WT Veh n= 21, WT Sob-AM2 n= 20, 5xFAD Veh n= 22, 5xFAD Sob-AM2 n= 21, F (3,83) =71.38. Cortex: WT Veh n= 21, WT Sob-AM2 n= 20, 5xFAD Veh n= 23, 5xFAD Sob-AM2 n= 22, F (3,82) =51.9.

### 3.8. Sob-AM2 increases phagocytic microglia in the hippocampus, but not cortex, of 5xFAD mice

The activation of phagocytic microglia in both the hippocampus and the cortex was measured by CD68 protein expression. In both regions, WT mice had minimal CD68 expression compared to 5xFAD mice, regardless of treatment (Figure 11A). Sob-AM2 treatment significantly increased CD68 expression in the hippocampus (Figure 11B) but not the cortex (Figure 11C) of 5xFAD mice.

**Figure 11:**
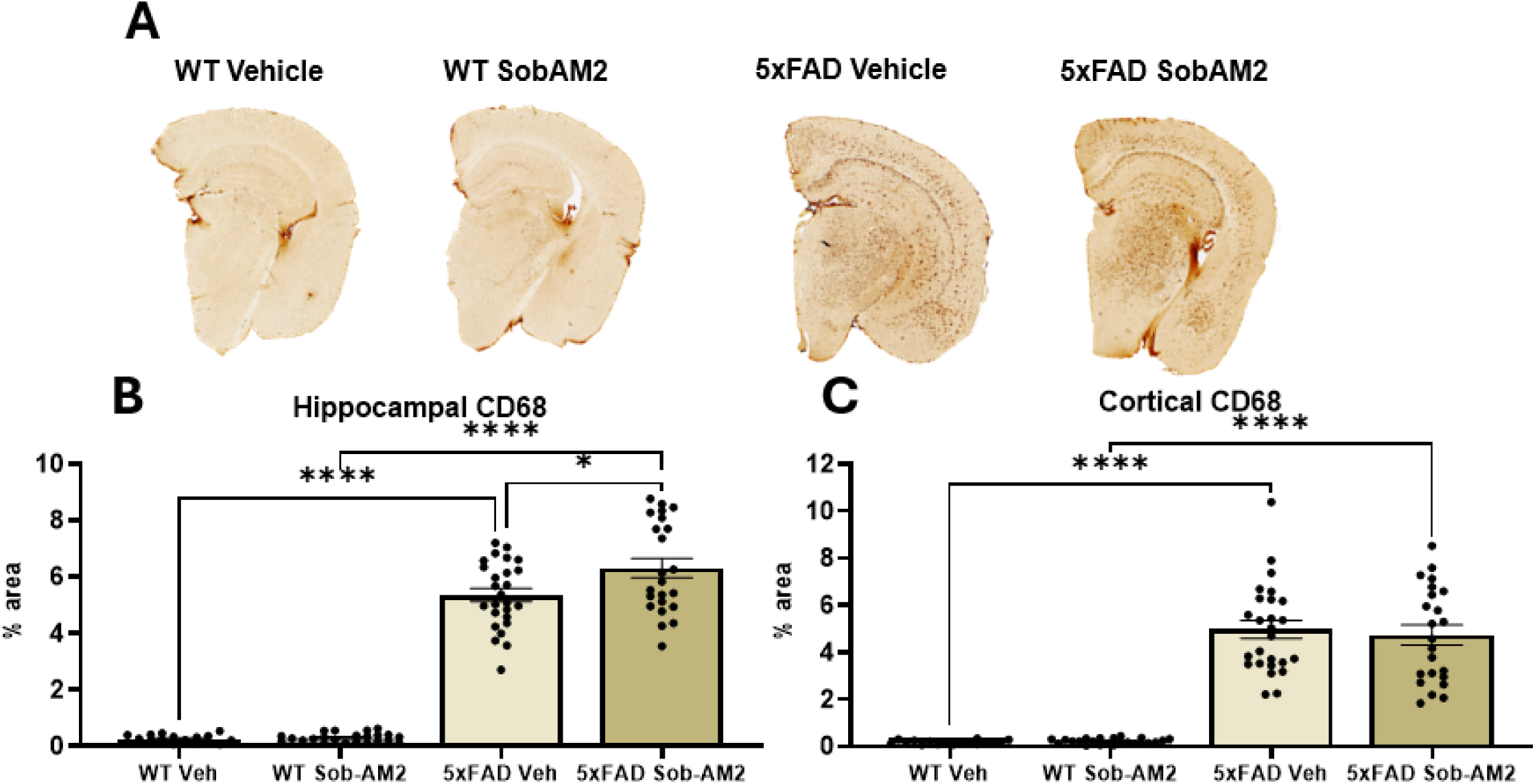
Sob-AM2 increases phagocytic microglia in the hippocampus, but not cortex, of 5xFAD mice. (A) Representative images of CD68 staining in WT and 5xFAD mice. 5xFAD mice displayed significantly higher CD68 expression compared to their WT littermate controls. Sob-AM2 treatment significantly increased CD68 expression in the hippocampus (B) of 5xFAD animals but not in the (C) cortex. *p<0.05, ****p<0.0001, Hippocampus: WT Veh n= 17, WT Sob-AM2 n= 23, 5xFAD Veh n= 25, 5xFAD Sob-AM2 n= 23, F (3,85) =204.7. Cortex: WT Veh n= 11, WT Sob-AM2 n= 17, 5xFAD Veh n= 25, 5xFAD Sob-AM2 n= 22, F (3,85) =74.72.

## 4. Discussion

This study sought to determine the effects of the TREM2-activating thyromimetic compound Sob-AM2 on cognition, Aβ pathology, and neuroinflammation in the 5xFAD mouse model of Aβ accumulation. Our novel dosing paradigm of twelve weeks of subcutaneous administration of Sob-AM2 at 100 ug/kg increased TREM2 expression, improved spatial and associative memory, increased synaptic density, and elevated phagocytic microglia in the hippocampus, without altering Aβ plaque burden or overall microglial activation. Sob-AM2 treatment did not alter body weight or overall mobility, suggesting a favorable safety profile.

The increased brain TREM2 protein concentration in Sob-AM2-treated mice confirms successful target engagement in the therapeutic target tissue. This was accompanied by the attenuation of memory deficits in the 5xFAD mice. These findings are consistent with previous reports of the effects of modulating TREM2 expression and activity in mice [15,26,33–36]. Treatment with a TREM2-agonizing antibody has been shown to significantly improve spatial, recognition, and contextual memory in 5xFAD mice [22, 30–32]. Similarly, TREM2 overexpression has significantly improved spatial and contextual memory in Aβ-accumulating mice [15, 23], while TREM2 knock-down worsened spatial memory [33]. Taken together, these findings appear to validate the therapeutic strategy of targeting TREM2 as a viable means of rescuing AD-related cognitive deficits.

In addition to the behavioral effects, we observed an increase in synaptic gene expression in Sob-AM2 treated 5xFAD mice. This aligns with previous reports of the effect of upregulating TREM2 expression and activity.

Both TREM2 overexpression and treatment with a TREM2-agonizing antibody have yielded significant increases in the expression of synaptophysin and PSD-95 as well as other synaptic genes [15, 23, 30, 34]. Interestingly, Sob-AM2 elicited a similar increase in synaptic gene expression in WT animals. The fact that this occurred without the accompanying improvement in CFR and OLM performance is surprising but perhaps reflects limitations of the particular behavioral assays. It is possible that WT animals were already performing so well in these tests that it was not possible to detect an appreciable improvement in the Sob-AM2 treated animals. Future studies incorporating more sensitive cognitive tests could help clarify this point.

There was no effect observed of Sob-AM2 on Aβ plaque in the 5xFAD mice. This is consistent with the results of a recent phase II trial of a TREM2 agonist, which reported that there was no decrease in plaque pathology in early-stage AD [35]. However, it has also been reported that people with missense TREM2 mutations, most notably the R47H variant, have an increase in the Aβ plaque burden of post-mortem human tissue [12, 36–38]. The literature on whether TREM2-targeting interventions promote Aβ clearance is mixed. Some studies using 5xFAD mice saw decreases in Aβ plaque burden resulting from a similar duration of treatment with a TREM2- agonizing antibody, as measured by immunohistochemistry [22, 30], while other studies using TREM2- agonizing antibodies in tau-seeded 5xFAD mice and PS2APP mice have not reported any alterations in plaque burden [39, 40]. This discrepancy may be related to the particular antibodies used to detect Aβ in each study.

The former two studies used a 6E10 antibody and a polyclonal antibody directed against Aβ residues 1-16, respectively. The 6E10 antibody is known to be cross-reactive with neuritic dystrophy marker APP [41], while the polyclonal antibody does not appear to have been validated for a lack of cross-reactivity. However, the latter two studies used the 3D6 and HJ3.4 antibody, both of which have been validated as being specific to Aβ [42, 43]. Our study used the MOAB-2 antibody, which has also been validated for its Aβ specificity [41]. Future studies that quantify both APP and Aβ are needed to clarify the effect of modulating TREM2 expression and activity on plaque pathology and neuritic dystrophy.

Sob-AM2 mediating an improvement in cognition without altering Aβ plaque burden could suggest an effect of the drug on soluble Aβ oligomer aggregation, rather than overt plaque pathology. Soluble Aβ oligomers are considered a stronger predictor of cognitive dysfunction than insoluble plaque accumulation [44]. This is particularly relevant in the context of therapeutically targeting TREM2, because while TREM2 can interact with a variety of Aβ isoforms, its highest affinity is with soluble Aβ oligomers [45]. This finding provides further motivation for future studies on TREM2 modulators to distinguish changes in specific Aβ isoforms as well as APP, which may provide a more precise understanding of the neurobiological processes underlying the cognitive benefits that many of these treatments appear to yield.

The fact that there was no observed effect of Sob-AM2 treatment on Iba1 expression contrasts with much of the literature on TREM2 modulation in Aβ mouse models. Chronic administration of TREM2-agonizing antibodies, which directly bind to the microglial TREM2 receptor, have tended to increase Iba1 expression in mouse models of Aβ accumulation [30, 31, 46]. Similarly, Aβ-overexpressing mice with a TREM2 deficiency or knock-out have shown a decrease in Iba1 expression [47–50], while TREM2 overexpression increased Iba1 immunostaining [34]. However, it should be noted that most of these studies assessed Iba1 immunostaining specifically within the area surrounding Aβ plaques rather than globally in different brain regions. Given that Iba1-positive microglia often cluster around Aβ plaques, this approach may yield more sensitive detection of subtle changes in microglial activation.

However, the lack of Sob-AM2-mediated microglial activation is not necessarily surprising given previous reports. In the spinal cords of EAE mice, Sob-AM2 significantly increased TREM2 expression without altering staining for another marker for microglial activation, CD11b [16]. The difference between Sob-AM2 and TREM2-agonizing antibodies in terms of microglial activation may stem from the fact that the TREM2- agonizing antibodies directly bind to TREM2 to activate the receptor while Sob-AM2 transcriptionally increases TREM2 expression [16]. It is possible that this transcriptional modulation of TREM2 expression may increase receptor availability on active microglia rather increasing TREM2 signaling. Analysis of downstream TREM2 targets could help clarify this issue. It is also possible that Sob-AM2 is affecting other microglial phenotypes beyond Iba1 expression. Future work, including a more in-depth analysis of microglial gene and protein expression, would allow for detection of more subtle shifts in microglial phenotypes.

In this study, there was an increase in hippocampal CD68 expression in response to Sob-AM2, but interestingly no effect was seen in the cortex. This is in contrast with several reports of TREM2 agonists resulting in increased CD68 expression in both the cortex as well as the hippocampus of 5xFAD mice [30, 31, 51]. However, it should be noted that these studies assessed CD68 staining specifically in the area surrounding Aβ plaques rather than globally in each region. Interestingly, TREM2-agonizing antibodies have not been reported to affect CD68 expression [40]. Still, this finding was not commensurate with the increase in CD68 we observed in the hippocampus.

One possible explanation for the regional specificity of the effect of Sob-AM2 on CD68 could be related to the thyromimetic effects of Sob-AM2. Hypothyroidism, a deficiency in thyroid hormones which may be an AD risk factor [52], has been shown to suppress microglial phagocytic activity [3] and AD-related hypothyroidism is regionally heterogeneous. In 5xFAD mice also expressing mutated human tau, the expression of downstream gene targets of T3 were decreased in the hippocampus but not the cortex [3]. In light of this finding, it is possible that administration of Sob-AM2, which mimics T3, may have a greater impact on the hippocampus than the cortex of murine AD models due to the differential impact of thyroid hormone signaling in those regions. In future studies, thyroid hormone signaling can be evaluated in different brain regions in response to Sob-AM2 treatment to evaluate whether this may account for the difference in CD68 expression resulting from SobAM2 administration.

## 5. Limitations

Some limitations should be considered. First the study only investigated a single dose of Sob-AM2 administered for a fixed duration. Evaluating an expanded dose response or optimizing timing of administration might impact the magnitude of responses observed. Second, this study utilized subcutaneous injections three times per week, which may have introduced additional variability. Repeated handling and injections could have increased stress, potentially influencing behavior outcomes, although these conditions were consistent across all treatment conditions. Third, some mice were singly housed due to fighting and aggressive behaviors.

Differences in social housing conditions may have affected sociability and cognitive performance, potentially influencing results obtained from behavioral assays. A final limitation is that we did not have data from all animals for all outcomes. In some cases, this was due to technical issues with the assays and in other cases due to non-participation of the animals. This missing data was distributed across all treatment conditions, but it nonetheless reduced our overall sample size for some outcomes and may have diminished our ability to detect more subtle changes.

## 6. Conclusions

The results of this study provide encouraging evidence for the therapeutic potential of SobAM2 to attenuate AD-mediated cognitive decline. Ongoing and future studies will enable us to determine the optimal timing of SobAM2 intervention and assess its impacts in other AD mouse model. Further research is needed to clarify the biological mechanisms through which the SobAM2-induced increase in TREM2 expression is enabling these cognitive improvements and how these may differ from the mechanisms associated with direct TREM2 activation.

## List of abbreviations

Aβ: amyloid-beta
AD: Alzheimer’s disease
APP: amyloid precursor protein
BCA: bicinchoninic acid
CFR: Conditioned Fear Response
CNS: central nervous system DAB: diaminobenzidine
DAM: disease-associated microglia
EAE: experimental autoimmune encephalomyelitis
IACUC: Institutional Animal Care and Use Committee
NMR: nuclear magnetic resonance
OF: Open Field
OLM: Object Location Memory
PBS: phosphate-buffered saline
PS1: presenilin 1
THR: thyroid hormone receptor
TREM2: Triggering Receptor Expressed on Myeloid Cells 2
T3: triiodothyronine
WT: wild-type

## Declarations

None

## Author Contributions

Investigation: L.K., G.A.J., W.H., S.V., N.G.K., F.K., S.K.; Data curation: L.K, G.A.J; Visualization: L.K., G.A.J, N.E.G.; Resources: T.B.; Conceptualization: T.S.S., J.F.Q., N.E.G.; Formal analysis: L.K., G.A.J., J.F.Q, N.E.G.; Funding acquisition: N.E.G.; Writing- original draft: L.K., G.A.J.; Writing – review and editing: L.K., G.A.J., T.S.S., J.F.Q., N.E.G.. All authors read and approved of the final manuscript.

## Institutional Review Board Statement

The animal work in this study was conducted in accordance with the NIH Guidelines for the Care and Use of Laboratory Animals and was approved by the institutional Animal Care and Use Committee of the Portland VA Healthcare System (ACORP #6155-23).

## Informed Consent Statement

Not applicable.

## Data Availability Statement

The datasets used and/or analyzed in the current study are available from the corresponding author upon reasonable request.

## Acknowledgments

Not applicable.

