## Supplemental Figures for "The TREM2 targeting small molecule Sob-AM2 improves cognition independent of Aβ plaque alteration in 5xFAD mice"

**Supplementary Figures**

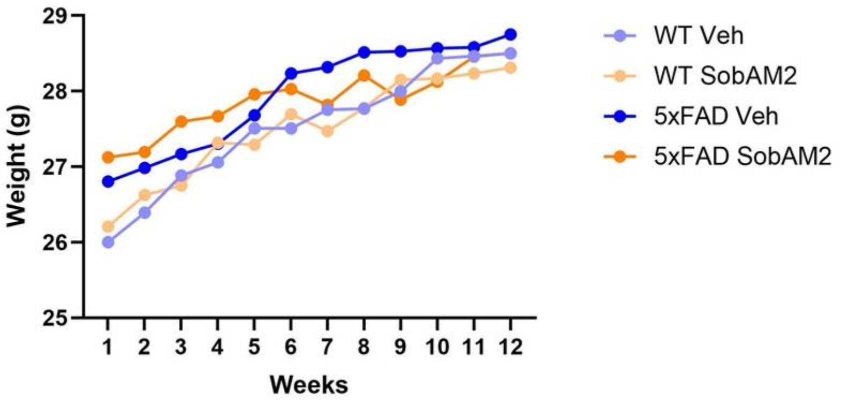


**Supplementary Figure 1: Body weight over the course of 12 weeks of treatment.**

| Left Fore | | Swing | Brake | Propel | Stance | Stride | Stance/Swing | Stride Length | Stride Frequency | Paw Angle | *Absolute Paw Angle | Paw Angle Variability | Stance Width | Step Angle | SLVar | SWVar | Step Angle Var |
|---|---|---|---|---|---|---|---|---|---|---|---|---|---|---|---|---|---|
| WT | Veh | 0.115 | 0.075 | 0.083 | 0.158 | 0.273 | 1.447 | 5.447 | 3.747 | -6.226 | 8.437 | 7.284 | 1.832 | 63.811 | 1.071 | 0.391 | 14.744 |
| | SobAM2 | 0.094 | 0.084 | 0.082 | 0.165 | 0.259 | 1.809 | 5.173 | 3.955 | -4.714 | 5.395 | 7.041 | 1.786 | 63.236 | 1.225 | 0.438 | 15.054 |
| 5xFAD | Veh | 0.093 | 0.075 | 0.076 | 0.151 | 0.244 | 1.741 | 4.859 | 4.288 | -2.112 | 6.312 | 8.800 | 1.841 | 66.000 | 1.222 | 0.492 | 15.757 |
| | SobAM2 | 0.097 | 0.079 | 0.077 | 0.156 | 0.253 | 1.671 | 5.057 | 4.062 | -3.605 | 5.729 | 7.738 | 1.738 | 65.886 | 1.161 | 0.402 | 13.998 |
| Left Hind | | | | | | | | | | | | | | | | | |
| WT | Veh | 0.084 | 0.069 | 0.110 | 0.179 | 0.263 | 2.147 | 5.253 | 3.895 | -19.300 | 19.300 | 5.542 | 2.895 | 51.205 | 0.863 | 0.270 | 14.261 |
| | SobAM2 | 0.085 | 0.067 | 0.111 | 0.179 | 0.264 | 2.155 | 5.286 | 3.891 | -15.986 | 15.986 | 5.818 | 2.891 | 50.268 | 0.927 | 0.287 | 14.506 |
| 5xFAD | Veh | 0.088 | 0.066 | 0.105 | 0.171 | 0.259 | 1.982 | 5.176 | 4.000 | -15.253 | 15.253 | 6.159 | 2.994 | 53.100 | 0.859 | 0.282 | 14.749 |
| | SobAM2 | 0.088 | 0.065 | 0.110 | 0.176 | 0.264 | 2.014 | 5.290 | 3.871 | -19.952 | 19.952* | 5.129 | 2.900 | 51.100 | 0.873 | 0.267 | 14.174 |
| Right Fore | | | | | | | | | | | | | | | | | |
| WT | Veh | 0.107 | 0.080 | 0.074 | 0.154 | 0.261 | 1.500 | 5.221 | 3.937 | 8.542 | 9.374 | 7.726 | - | - | 1.041 | - | - |
| | SobAM2 | 0.094 | 0.089 | 0.080 | 0.169 | 0.262 | 1.832 | 5.255 | 3.918 | 4.877 | 7.205 | 8.405 | - | - | 1.132 | - | - |
| 5xFAD | Veh | 0.095 | 0.089 | 0.073 | 0.162 | 0.257 | 1.824 | 5.129 | 4.053 | 4.606 | 7.724 | 7.965 | - | - | 1.271 | - | - |
| | SobAM2 | 0.100 | 0.083 | 0.075 | 0.158 | 0.258 | 1.610 | 5.162 | 3.971 | 6.433 | 8.529 | 8.281 | - | - | 1.307 | - | - |
| Right Hind | | | | | | | | | | | | | | | | | |
| WT | Veh | 0.083 | 0.066 | 0.110 | 0.177 | 0.259 | 2.168 | 5.200 | 3.937 | 19.379 | 19.379 | 5.537 | - | - | 0.864 | - | - |
| | SobAM2 | 0.085 | 0.074 | 0.106 | 0.180 | 0.264 | 2.214 | 5.286 | 3.891 | 18.677 | 18.677 | 5.882 | - | - | 0.930 | - | - |
| 5xFAD | Veh | 0.086 | 0.067 | 0.098 | 0.165 | 0.250 | 1.982 | 5.006 | 4.247 | 16.276 | 16.276 | 5.676 | - | - | 0.884 | - | - |
| | SobAM2 | 0.086 | 0.071 | 0.106 | 0.178 | 0.264 | 2.110 | 5.281 | 3.895 | 19.067 | 19.067 | 5.652 | - | - | 0.928 | - | - |

| Left Fore | | #Steps | Stride Length CV | Stance Width CV | Step Angle CV | Swing Duration CV | Paw Area | Paw Area Variability | Stance Factor | Gait Symmetry | MAX dA/dT | MIN dA/dT | Overlap Distance | PPP | Ataxia Coefficient | Midline Distance | Axis Distance |
|---|---|---|---|---|---|---|---|---|---|---|---|---|---|---|---|---|---|
| WT | Veh | 36.526 | 19.811 | 21.429 | 23.759 | 25.935 | 0.316 | 0.038 | 0.987 | 1.015 | 19.704 | -6.678 | 1.017 | 0.532 | 0.932 | -2.415 | -0.889 |
| | SobAM2 | 44.773 | 23.654 | 25.041 | 24.399 | 28.756 | 0.370 | 0.064 | 0.999 | 1.004 | 24.902 | -8.252 | 1.285 | 0.525 | 1.186 | -2.081 | -0.855 |
| 5xFAD | Veh | 31.912 | 25.609 | 27.545 | 24.859 | 27.711 | 0.378 | 0.063 | 0.999 | 1.011 | 26.349 | -8.810 | 0.777 | 0.531 | 1.158 | -2.271 | -0.945 |
| | SobAM2 | 38.905 | 23.388 | 23.665 | 21.861 | 29.177 | 0.359 | 0.054 | 0.961 | 1.014 | 25.240 | -7.326 | 1.472 | 0.573 | 1.096 | -2.250 | -0.849 |
| Left Hind | | | | | | | | | | | | | | | | | |
| WT | Veh | 41.079 | 16.483 | 9.568 | 28.152 | 27.115 | 0.698 | 0.089 | 1.012 | 1.015 | 60.454 | -12.571 | 1.447 | 0.599 | 0.820 | 2.044 | -1.499 |
| | SobAM2 | 35.500 | 17.724 | 10.047 | 28.923 | 25.795 | 0.785 | 0.099 | 0.996 | 1.010 | 69.224 | -13.782 | 0.905 | 0.476 | 0.975 | 2.081 | -1.456 |
| 5xFAD | Veh | 40.147 | 16.916 | 9.472 | 29.299 | 22.683 | 0.818 | 0.110 | 1.055 | 0.995 | 72.901 | -16.084 | 1.209 | 0.551 | 0.896 | 2.199 | -1.512 |
| | SobAM2 | 39.119 | 16.800 | 9.223 | 27.922 | 28.445 | 0.738 | 0.080 | 0.989 | 1.021 | 66.122 | -14.240 | 1.130 | 0.549 | 0.786 | 2.000 | -1.521 |
| Right Fore | | | | | | | | | | | | | | | | | |
| WT | Veh | 40.421 | 20.223 | - | - | 28.807 | 0.298 | 0.053 | - | 1.015 | 19.209 | -6.548 | 1.426 | 0.496 | 1.003 | -2.169 | 0.912 |
| | SobAM2 | 34.750 | 21.831 | - | - | 30.230 | 0.333 | 0.061 | - | 1.010 | 21.884 | -6.697 | 1.100 | 0.668 | 0.975 | -2.039 | 0.851 |
| 5xFAD | Veh | 38.765 | 25.532 | - | - | 29.429 | 0.396 | 0.071 | - | 0.995 | 27.292 | -10.539 | 1.013 | 0.538 | 1.176 | -2.038 | 0.953 |
| | SobAM2 | 37.690 | 25.519 | - | - | 30.521 | 0.312 | 0.056 | - | 1.021 | 20.577 | -7.355 | 0.024 | 0.595 | 1.226 | -2.115 | 0.895 |
| Right Hind | | | | | | | | | | | | | | | | | |
| WT | Veh | 41.447 | 16.803 | - | - | 27.668 | 0.670 | 0.084 | - | 1.015 | 57.016 | -12.548 | 1.426 | 0.496 | 0.908 | 2.090 | 1.517 |
| | SobAM2 | 35.273 | 17.603 | - | - | 25.860 | 0.772 | 0.103 | - | 1.010 | 66.013 | -14.123 | 1.100 | 0.668 | 0.881 | 2.208 | 1.470 |
| 5xFAD | Veh | 41.529 | 19.043 | - | - | 28.509 | 0.825 | 0.101 | - | 0.995 | 71.299 | -17.573 | 1.013 | 0.538 | 0.957 | 2.201 | 1.529 |
| | SobAM2 | 38.714 | 17.730 | - | - | 26.445 | 0.762 | 0.087 | - | 1.021 | 65.304 | -14.580 | 0.024 | 0.595 | 0.874 | 2.078 | 1.474 |

**Table 1: DigiGate 32 Metric analysis of gait and coordination**. Averages represented for each treatment group and each metric. *$p<0.05$ in post-hoc pairwise comparison on 5xFAD SobAM2 vs 5xFAD vehicle, PPP= Paw Placement Positioning
